# Properties and computational consequences of fast dendritic spikes during natural behavior

**DOI:** 10.1101/2021.12.27.474234

**Authors:** Alain Destexhe, Mayank Mehta

## Abstract

Dendritic membrane potential was recently measured for the first time in drug-free, naturally behaving rats over several days. These showed that neuronal dendrites generate a lot of sodium spikes, up to ten times as many as the somatic spikes. These key experimental findings are reviewed here, along with a discussion of computational models, and computational consequences of such intense spike traffic in dendrites. We overview the experimental techniques that enabled these measurements as well as a variety of models, ranging from conceptual models to detailed biophysical models. The biophysical models suggest that the intense dendritic spiking activity can arise from the biophysical properties of the dendritic voltage-dependent and synaptic ion channels, and delineate some computational consequences of fast dendritic spike activity. One remarkable aspect is that in the model, with fast dendritic spikes, the efficacy of synaptic strength in terms of driving the somatic activity is much less dependent on the position of the synapse in dendrites. This property suggests that fast dendritic spikes is a way to confer to neurons the possibility to grow complex dendritic trees with little computational loss for the distal most synapses, and thus form very complex networks with high density of connections, such as typically in the human brain. Another important consequence is that dendritically localized spikes can allow simultaneous but different computations on different dendritic branches, thereby greatly increasing the computational capacity and complexity of neuronal networks.

## Introduction

Microelectrodes have been used for nearly a century to do long-term measurements of extracellular somatic action potentials in freely behaving subjects. However, somatic action potentials occur rarely (~1.5 Hz in rodent neocortical principal neurons) (Buzsáki & Mizuseki, 2014; Moore *et al.*, 2017) and do not provide an estimate of the intracellular dynamics. While the vast majority of excitatory synapses are localized on their dendritic arbors, spanning more than 1000 μm, most pyramidal neurons receive inhibition predominantly near the soma which can decouple the somatic and dendritic activities. Dendrites in vitro can generate local dendritic action potentials (DAP) (Spencer & Kandel 1960, Amitai et al. 1993, Nevian et al. 2007, Johnston & Narayanan 2008). Some back-propagating action potentials (bAP) initiated at cell somata can also reach the proximal but not the distal dendrites due to dendritic attenuation (Waters & Helmchen 2004) and inhibition. Two-photon calcium imaging can be used to study dendrites in vivo, but these can only study calcium dynamics, whose link with sodium spiking is not straightforward, and this does not allow a direct measure of subthreshold membrane potential or dendritic sodium spikes. Sharp electrode and patch-clamp techniques damage or rupture the membrane, thus limiting the recording duration (Deweese 2017) and altering in vivo neural dynamics including firing rates. Additionally, these methods often require the subject to be anesthetized or immobilized, which alter neural dynamics and limit possible behavioral paradigms.

Extensive experimental and computational studies have investigated the contribution of both the passive and active properties of dendritic properties on neural dynamics and learning. For example, dendritic depolarization can not only influence synaptic plasticity (Golding *et al.*, 2002) but also the propagation of sodium spikes (Waters & Helmchen, 2004). This can influence the organization of synapses and confer local computational capacity to dendrites (Rall & Segev, 1988; Mel, 1993; Poirazi & Mel, 2001; Mehta, 2004). We review some exciting experimental and theoretical developments in these directions.

### Consequences of excitation inhibition balance and dendritic nonlinearities

Excitation and inhibition need to be in balanced for normal brain function. But, neurons are extended objects with asymmetric distribution of excitation and inhibition –cortical dendrites contain all the excitatory synapses, while inhibition is mostly focused near the soma. There are nearly five times as many excitatory synapses as inhibitory on dendrites. The excitation inhibition balance at the level of neurons and networks therefore means that there would be a major excitation inhibition imbalance within a neuron with excessive excitation in the distal dendrites and excessive inhibition at the soma and proximal dendrites. Additionally, dendrites have plenty of active channels, including voltage gated sodium and potassium channels. These commonly accepted ideas make a radical prediction –the distal dendrites could generate a lot of dendritic sodium spikes from local excitation, but the majority of them would be fail to generate somatic spike due to perisomatic inhibition. Somatic spike propagation in the dendrites too would be attenuated, causing further decoupling of soma and dendrites.

This would profoundly impact neural coding, computation and plasticity. For example, classic in vitro studies showed that the NMDAR-dependent synaptic plasticity can be induced on distal dendrites via the locally generated depolarization or dendritic spikes, in the complete absence of somatic spiking(Golding *et al.*, 2002). Theoretical studies suggested that this would result in a novel, dendritically constrained Hebbian learning rule and a novel excitation-inhibition synaptic cluster of fundamental memory unit (Mehta, 2004) (Figure 1).

**Figure 1.**
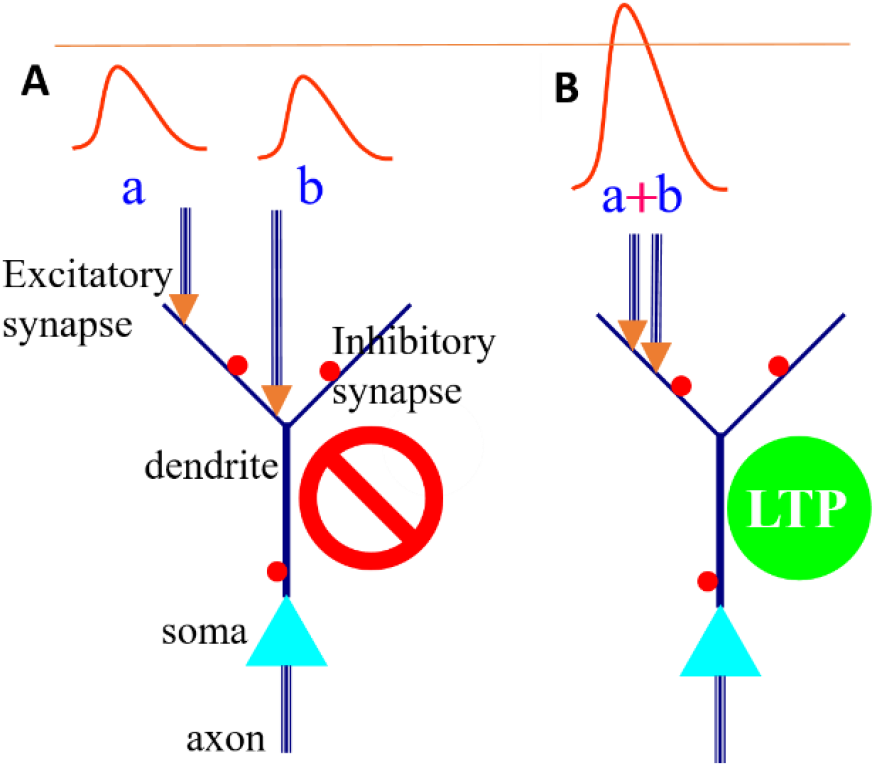
Cooperative long-term potentiation (LTP) can map similar memories on nearby locations on the dendrite. **A** Glutamatergic synapses (blue arrows) from two inputs (i.e. groups of neurons) a and b are located on different dendritic branches of neuron C. Red dots represent inhibitory synapses. Excitatory postsynaptic potentials (EPSPs) resulting from coactivation of a and b would be decremented or even be eliminated by an inhibitory synapse located between the two neurons before they could coalesce to generate a large dendritic spike or LTP. The dendritic distance-dependent attenuation of voltages would further curtail cooperation between input from different dendrites. **B** The two excitatory synapses are near each other, so their EPSPs could summate without being affected by dissipation or inhibition, to cross sodium spike threshold (thin blue line) and trigger a dendritic spike, whose large depolarization is sufficient to remove the magnesium block and induce LTP. Thus, the information about coactivation of events represented by a and b would be more likely to be encoded by nearby synapses (B) than dispersed synapses (A). Thus dendritically constrained Hebbian learning rule states that: “Synapses from neurons A and B onto a postsynaptic neuron C will be potentiated if neurons A and B are repeatedly coactivated and if their synapses are located near each other on a dendrite of neuron C”. The fundamental unit of memory would be a local cluster of excitatory synapse flanked by inhibitory synapses (Adapted from (Mehta, 2004)).

This has profound consequences, including clustering of synapses representing temporally similar information, and dendritic branch specific computations. However, testing this has remained a challenge since it has not been possible to measure dendritic depolarization and spiking from the thin dendrites in freely behaving animals until recently. In fact, conditions that generate dendritic spikes in vitro, e.g. synchronous stimulation of a cluster of inputs on a dendrite and blocked inhibition, may not exist in vivo. Hence, several studies have questioned the possibility of generating dendritic spikes during natural behavior and the vast majority of computational models of neuronal networks do not include dendritic spiking.

Another set of biophysical theories of dendritic processing predicted that there would be an optimal frequency for inducing NMDAR-mediated plasticity at a single synaptic spine, such that not only the lower but even higher frequency stimuli would induce lesser plasticity (Kumar & Mehta, 2011). Further, the theory predicted that the preferred frequency for plasticity would increase with increasing distance of the spine from the soma when the plasticity relies on somatically generated back-propagating action potential (Kumar & Mehta 2006, 2007). Even more surprising, the theory predicted that periodic stimuli would induce far greater LTP and LTD, than aperiodic stimuli, at a single NMDAR-mediated synapse(Kumar & Mehta, 2011).

While several studies support these novel forms of dendritic computations, until recently it has not been possible to test these important yet surprising predictions and their consequences, due to technological limitations. The standard technique for measuring the somatic membrane potential is patch clamp. This works very well in vitro for the soma and proximal dendrites, that have a large diameter and are relatively stiff. But, the vast majority of excitatory synapses are located on thin dendritic branches, including apical, oblique and basal dendrites, which are difficult to patch. Recent advances have measured the dendritic membrane potential on the apical shaft in head fixed rodents up to 400um from the soma, but the activity of the thin branches and distal dendrites in vivo and even in vitro have remained a mystery. Calcium imaging has made great strides in measuring dendritic calcium spikes, these are related to sodium spikes but the relationship is not straightforward. Further, calcium imaging does not measure membrane potential. Developments of novel, voltage sensitive dyes may address these challenges. But, issues such as separating signal from background and measuring deeper brain regions’ activity would be a challenge.

### Dendritic membrane potential measurements during natural behavior

A novel approach was developed recently to overcome these challenges and measure the dendritic membrane potential in freely behaving rat for up to four days. The technique leveraged basic principles of electricity, namely a voltage divider circuit, to measure dendritic membrane potential without puncturing the dendrite (Moore et al. Science 2017, figure 1).

Briefly, patch-clamp technique can measure cellular membrane voltage because of the electrical resistance between the liquid in the pipette and the intracellular fluids is much smaller than the impedance between the pipette and the extracellular medium, resulting in current flow through the pipette. The novel technique involved a two-step process. The first step was of placing a flexible tetrode very near a dendrite. The flexible nature of the tetrode ensured that it moved with the surrounding brain, thus minimizing damage to the nearby dendritic processes. Keeping the electrode near the dendrite without puncturing it kept the dendrite intact allowing long-term measurement. The second step was to let the glial cells encapsulate the electrode and the nearby dendritic process. Such glial encapsulation occurs naturally, as a response to the small injury caused by the electrode. The large blob of glial tissues generates a large resistance between the dendrite-electrode assembly and ground and this resistance is much larger than the resistance between the thin membrane of a dendrite and the electrode. Thus, this voltage divider circuit allows one to measure the dendritic membrane potential, without puncturing the dendrite. The glial sheath holds together the dendrite-electrode assembly enabling stable recording for more than four days, a major step forward.

### Dendrites generate five times as many sodium spikes as the soma in all cortical layers

These experiments provided the first measurements of cortical dendritic membrane potential and spiking in freely behaving rats. The measurements were obtained not only from the most superficial cortical tissue but also deeper tissue, from 200um from the cortical surface to 1500um. Thus, these DAP arise from the neural somata in all layers and all parts of dendrites, including the apical most, thin dendrites, and the basal dendrites. The rise times of these DAP were quite short (about 0.5ms) and quite consistent across different DAP sources, clearly indicative of sodium spikes. But, the decay times and widths were quite variable, ranging from a few milliseconds to tens of milliseconds (Figure 4). Most of them were not accompanied by a calcium spikes, evidenced by DAP decay times less than 50ms. But, some of these sodium spikes could have triggered a calcium or NMDA spike, evidenced by longer decay times. The measured DAP amplitudes were quite variable, determined by the resistance of the glial seal. The highest measured DAP amplitude was 25mV.

**Figure 2:**
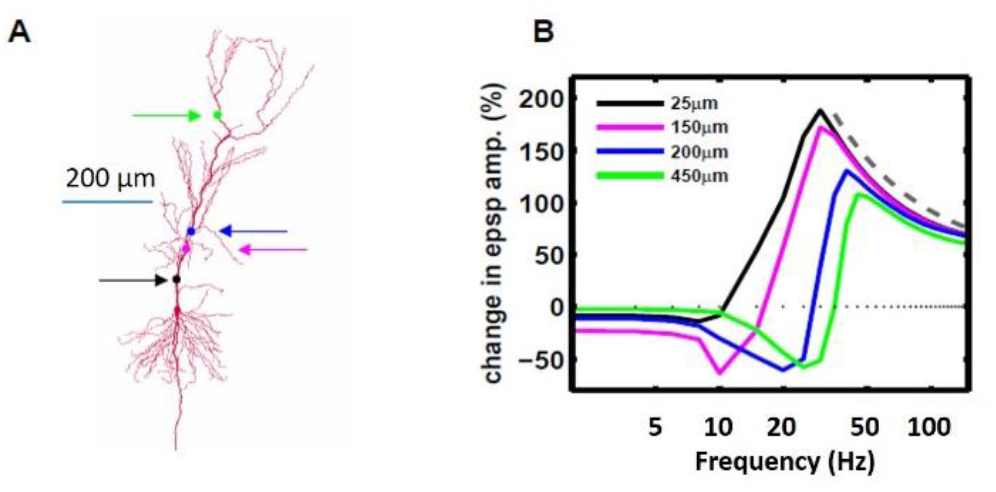
Biophysical model predicts dendritic location dependent optimal frequency for inducing NMDAR-dependent plasticity. **A**. A biophysical model of an excitatory neuron, arrows indicate the locations where the model NMDAR-dependent synaptic spine was placed. **B**. The magnitude of LTP as a function of coincident activation of NMDAR-synapse and postsynaptic somatic spiking. Only 10 spikes were generated to ensure LTP did not saturate. Notice that there is an optimal frequency for inducing maximal LTP, such that increasing frequency results in lesser LTP. The optimal frequency increases with increasing distance along the dendrite from the soma (Kumar & Mehta, 2006, 2007. 2009, Adapted from (Kumar & Mehta, 2011)).

**Figure 3:**
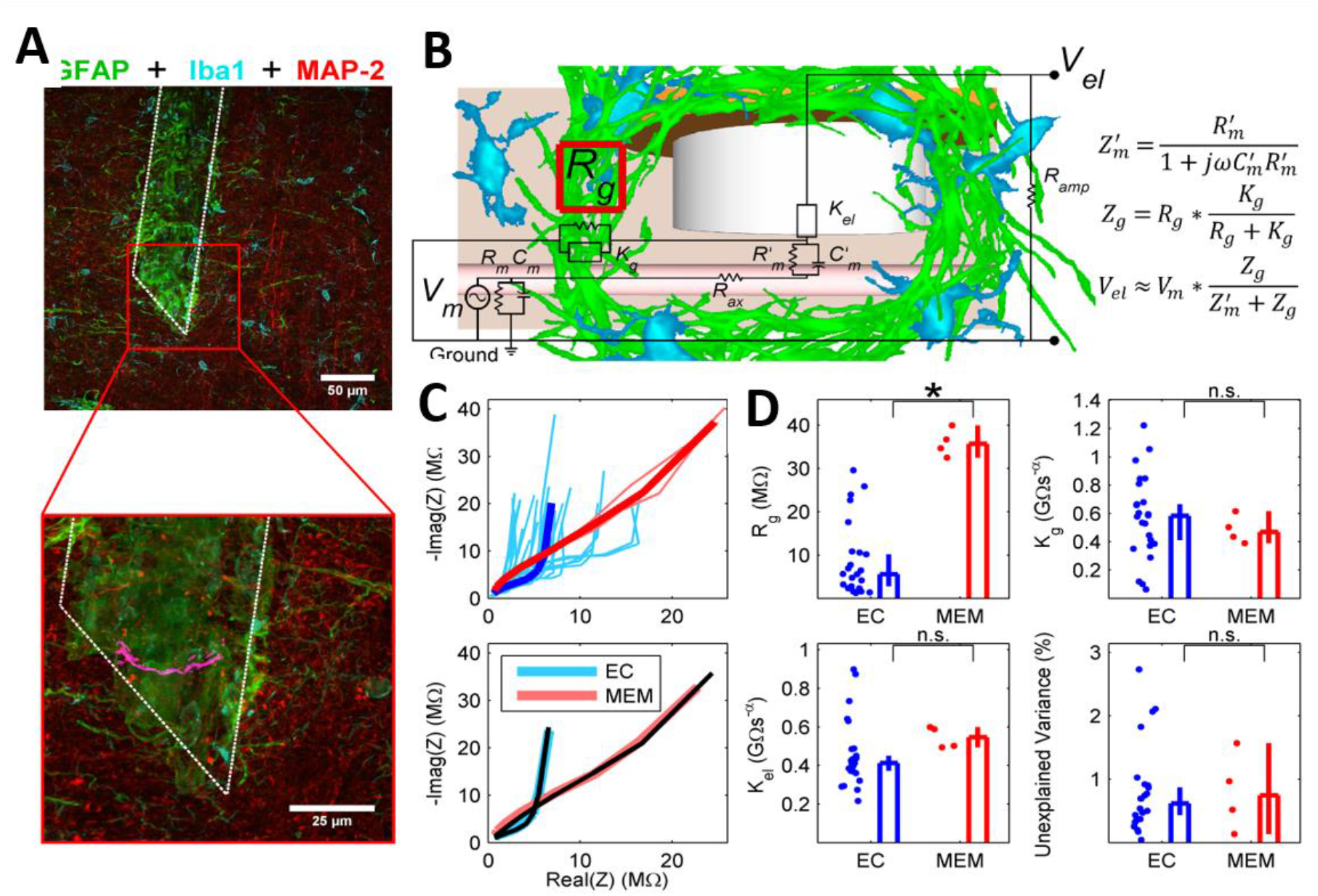
Glial sheath mechanism of DAP recordings. **A.** Top, maximum intensity projection of cortical slice immunohistologically labeled for GFAP (green), Iba1 (cyan), and MAP-2 (red) (see Methods). The white dotted line outlines the putative region where the tetrode was placed, ~650 μm from the pial surface of Posterior Parietal Cortex. The aggregation of GFAP-positive reactive astrocytes is evident encapsulating the tetrode site. Bottom, semi-transparent projection of tetrode tip region. An oblique and intact dendrite segment passing through the encapsulated area is highlighted in magenta. Due to the chronic nature of these experiments we can’t confirm that data were recorded from this dendrite, but its proximity to the encapsulated region illustrates the feasibility of our hypothesized mechanism. See Movie S2 for 3-dimensional semi-transparent projection. **B.** Equivalent electrical circuit. The voltage difference between the tetrode tip and ground is proportional to the voltage difference between the inside of the dendrite and ground; the magnitude depends on the relative values of the glial sheath impedance *Z*_*g*_ and membrane impedance *Z’*_*m*_. In typical extracellular recordings, *Z*_*g*_ is negligible compared to *Z’*_*m*_, so no intracellular signal is recorded. Here *j* is the imaginary number and ω is frequency. **C.** Top, impedance spectra for normal extracellular (EC, light blue) and DAP-recording (MEM, light red) electrodes shows increased impedance for the DAP-recording electrodes. Dark, bold lines represent the mean impedance spectra. Bottom, fitting the electric circuit model in (b) provides a close approximation (solid black lines) to the mean sample impedance for both extracellular (light blue line) and DAP-recording tetrodes (light red line). **D.** The fitted model parameter *R*_*g*_ (glial sheath resistance) for MEM (top left) was significantly larger than *R*_*g*_ for EC, consistent with our proposed glial sheath recording mechanism. In contrast, the fitted model parameters for *K*_*g*_ (top right, EC, MEM) and *K*_*el*_ (bottom left, EC, MEM) were not significantly different from each other. The percentage variance unexplained values (bottom right) for both EC and MEM were very low, and not significantly different from each other. (Moore & Mehta, 2012, 2013, 2014, Adapted from (Moore *et al.*, 2017)).

**Figure 4:**
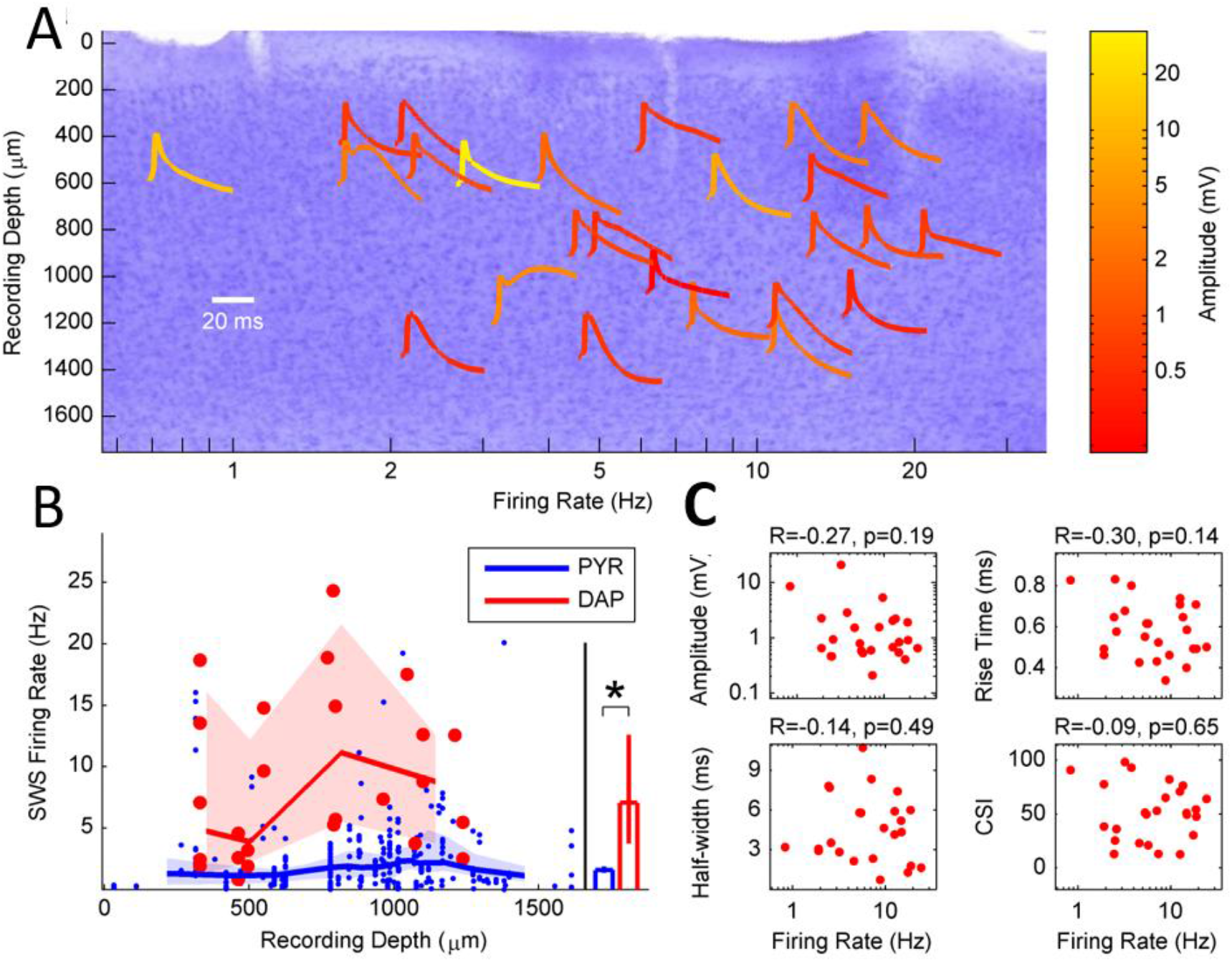
Depth independence of DAP waveform and rate. **A.** Background shows Nissl stained section of a representative parietal cortical tissue from which DAP were recorded. Tetrode tracks are seen are vertical white streaks. Different DAP waveforms, averaged across the respective session, recorded from all animals, are shown as a function of the recording depth and DAP rate. Scale bar shows the duration of the waveforms. Notice that only a handful of DAP waveforms are wide, indicative of calcium spike following the DAP. The majority of DAP are narrow, indicative of only sodium spiking. The DAP rate was not significantly correlated with recording depth. **B.** The average DAP rate was computed as a function of the recording depth across all data. This was compared with the average firing rates of ensembles of pyramidal neurons’ somatic spikes as a function of depth, measured under similar conditions as DAP. DAP rates were far greater than pyramidal neuron somatic spike rates at all depths. **C.** DAP mean firing rate was not significantly correlated with amplitude, rise time, half-width, or CSI, demonstrating the robust and unbiased nature of the findings (Moore & Mehta, SfN 2012, 2013, 2014, Adapted from (Moore *et al.*, 2017)).

Remarkably, the average DAP rates were very high, nearly five times the simultaneously measured somatic spiking rates. This can’t be explained by nonspecific variables such as the recording depth, the measured DAP amplitude, the DAP rise time, duration of recording (ranging from half an hour to four days), DAP decay time or width, or the activity-dependent adaptation of the DAP properties. The DAP recordings were remarkably stable across the long duration of measurements, showing no consistent drift. Thus, the very high DAP rates are unlikely to arise due to artefacts or the measurement process and likely a result of genuine, undisturbed mechanisms dendritic processing. Indeed, the DAP rates varied consistently, in a sigmoid fashion, as a function of the subthreshold dendritic membrane potential, with very small variability in DAP amplitude across the entire experiment. In fact, the DAP amplitude was always smaller than the range of fluctuations of the subthreshold dendritic membrane potential, which has not been seen in the somatic or proximal dendritic recordings. In fact, subthreshold dendritic membrane potential fluctuations exceeded 25mV in some measurements, and the true range of fluctuations were likely higher. These findings along with the large DAP rates from the somatic rates, demonstrate a major dissociation between dendritic processing from the somatic spiking.

Remarkably, while the DAP rates were five times greater than the somatic rates when the animals were naturally immobile, this disparity became even larger when the animals were ambulating. Both the somatic spike rates and DAP rates increased when the rats were walking around in a maze. But, the dap rates now became ten times greater than the somatic rates. These results demonstrate not only a decoupling of dendritic spiking from the somatic spiking, but a nonlinear amplification of synaptic inputs by dendritic active conductances, such that the amplification doubled during active exploration compared to resting.

### Biophysical models of fast dendritic spikes

We use here a detailed computational model of dendritic spiking activity that was designed in previous papers (Destexhe and Pare, 1999; Rudolph and Destexhe, 2003). The model was based on reconstructed morphologies from cat parietal cortex, in which neurons were recorded in vivo. This model puts a particular emphasis on reproducing in vivo electrophysiological measurements, obtained intracellularly (Pare et al., 1998). These measurements provided, for the first time, intracellular recordings of the same neurons during active states, and after total suppression of activity using extracellular application of tetrodotoxin (TTX). TTX suppressed all Na channel activity and therefore suppressed completely network activity inducing a flat local LFP, and revealing the natural resting membrane properties of the cell. These data were the first to provide quantitative measurements of the synaptic background activity in vivo, and were used to construct biophysical models, taking into account the actual densities of excitatory and inhibitory synaptic terminals in soma and dendrites of cat pyramidal neurons, and the quantal conductance at each synapse (see details in Destexhe and Pare, 1999).

The intracellular measurements comparing intact network activity with suppressed activity, revealed that synaptic background activity is responsible for a depolarization, an increase of conductance relative to rest, and an increase of Vm fluctuations (reviewed in Destexhe et al., 2003). Conductance measurements were later obtained from intracellular recordings in awake and sleeping cats (Rudolph et al., 2007), and confirmed the depolarization and Vm fluctuations, while the increase of conductance was more moderate. In different brain states in cats, such as wakefulness, slow-wave sleep up states, and paradoxical sleep, similar properties were observed. These measurements are still unique today, as there is no other characterization of neurons in the same brain area, in awake animals, and in the same cells before and after suppression of network activity.

The biophysical models based on such measurements revealed a number of interesting computational properties (reviewed in Destexhe et al. 2003). We focus here on the local effect of synaptic activity on dendritic dynamics, and delineate a series of consequences that may be important for the type of computations performed by dendrites in vivo.

#### Consequence 1: Synaptic background activity facilitates the initiation of fast dendritic spikes

Figure 5 shows the spiking activity of this biophysical model, comparing soma and dendrites. It is apparent that the dendrites generate an intense local dendritic spike activity, few of which reach the soma and lead to somatic spiking. This feature was constantly observed in these models, and is examined in more detail below.

**Figure 5:**
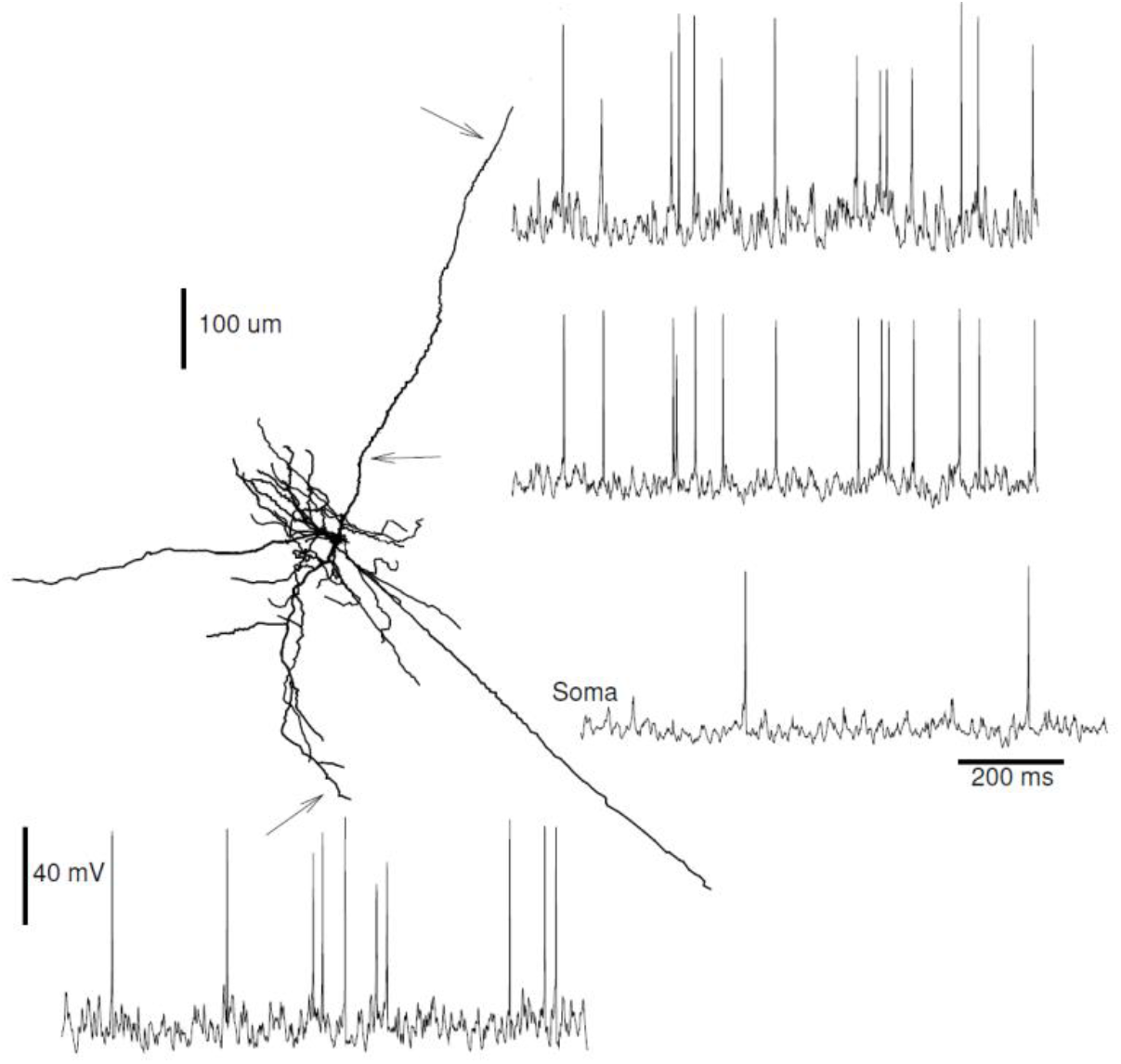
Intense local dendritic spike activity in the dendrites of a biophysical model of synaptic background activity in vivo. A biophysical model of a layer 6 pyramidal neuron from cat cortex was run with randomly releasing excitatory and inhibitory synapses at low rates, which was adjusted to mimic in vivo measurements. The activity of the model is shown in the soma, where spiking activity was relatively sparse, and in three dendritic locations (arrows), where the local dendritic spike activity was more intense. Model from Destexhe and Pare, 1999.

To explain this phenomenon of intense local dendritic spikes, the dynamics of the initiation and propagation of fast dendritic spikes was investigated. First, dendritic spike propagation was simulated in computational models of four morphologically-reconstructed cortical pyramidal neurons which included voltage-dependent currents in soma, dendrites and axon (see Methods in Destexhe and Pare, 1999). As shown in Fig. 6A (top), in quiescent conditions (no synaptic background activity), backpropagating dendritic spikes were reliable up to a few hundred microns from the soma (Fig. 6A, bottom, quiescent), in agreement with dual soma/dendrite recordings in vitro (Stuart et al., 1997). In the presence of synaptic background activity, backpropagating spikes were still robust, but propagated over a more limited distance in the apical dendrite compared to quiescent states (Fig. 6A, bottom, in-vivo–like), consistent with the limited backwards invasion of apical dendrites observed with two-photon imaging of cortical neurons in vivo (Svoboda et al., 1997). Thus, the model is consistent with backpropagating spikes, as typically observed in pyramidal cell dendrites.

**Figure 6:**
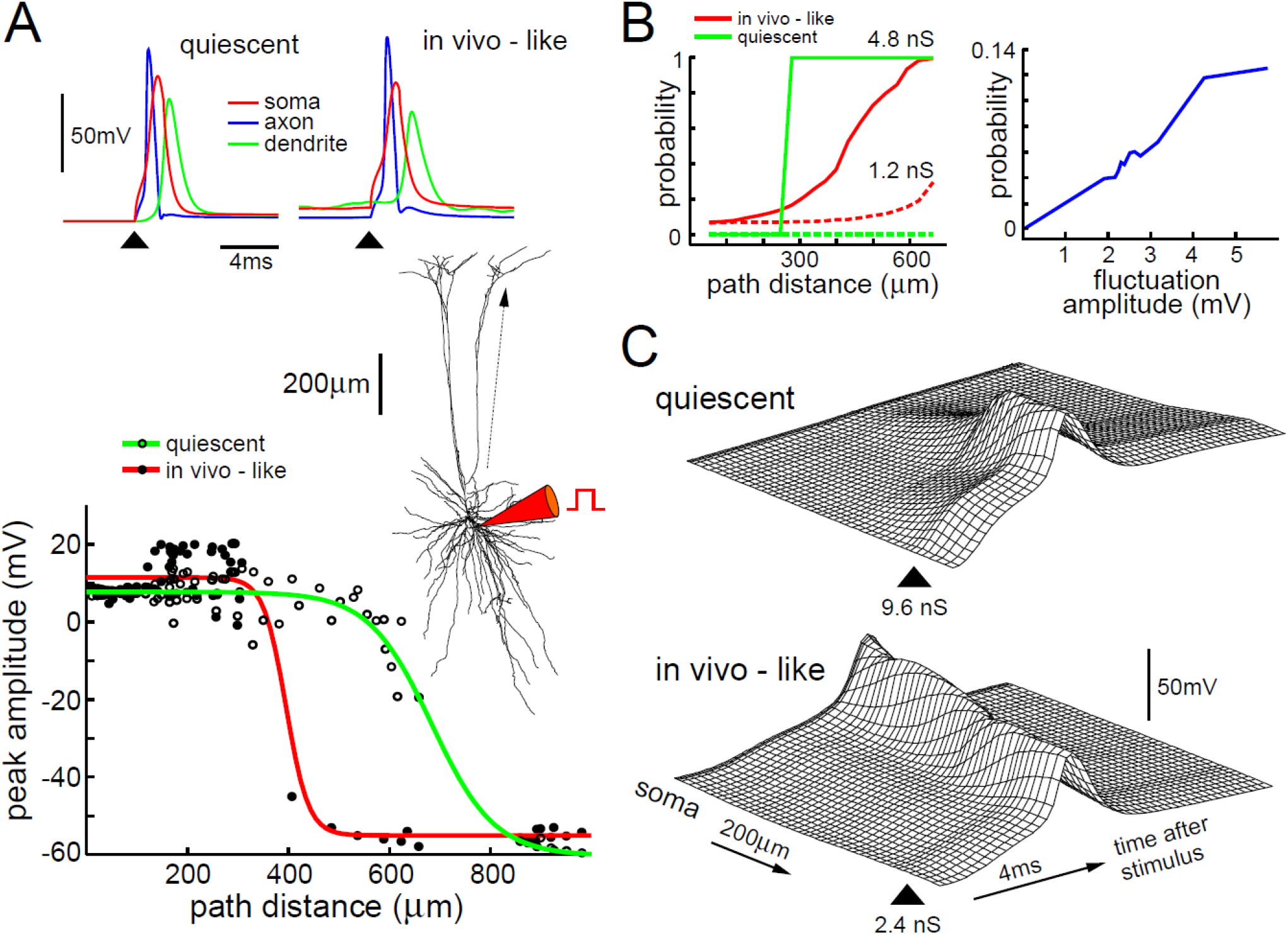
Initiation and propagation of fast dendritic spikes in a biophysical model of pyramidal cell under in vivo–like activity. **A**. Impact of background activity on spike backpropagation in a layer V cortical pyramidal neuron. Top: the respective timing of spikes in soma, dendrite (300 μm from soma) and axon are shown following somatic current injection (arrow). Bottom: spike backpropagation in the apical dendrite for quiescent (green) and in-vivo–like (red) conditions. The backwards invasion was more restricted in the latter case. **B**. Impact of background activity on the initiation of fast dendritic spikes. Left: probability for initiating a dendritic spike shown as a function of path distance from soma (left) for two different amplitudes of AMPA-mediated synaptic stimuli (dash: 1.2 nS; solid: 4.8 nS). Right: probability of dendritic spike initiation (100 μm from soma) as a function of the amplitude of voltage fluctuations (1.2 nS stimulus). **C**. Impact of background activity on dendritic spike propagation. A forward-propagating dendritic spike was evoked in a distal dendrite by an AMPA-mediated EPSP (arrow). Top: in quiescent conditions, the spike only propagated within 100-200 μm, even for high-amplitude stimuli (9.6 nS shown here). Bottom: under in-vivo–like conditions, dendritic spikes could propagate up to the soma, even for small stimulus amplitudes (2.4 nS shown here). The distance axis corresponds to the arrow indicated in the cell morphology in A. Modified from Rudolph and Destexhe, 2003.

Fast spikes could also be initiated in dendrites following simulated synaptic stimuli (Stuart et al., 1997). In quiescent conditions, the threshold for dendritic spike initiation was high (Fig. 6B, left, quiescent), and the dendritic-initiated spikes forward-propagated only over limited distances (100-200 μm; Fig. 6C, quiescent), in agreement with previous observations (Stuart et al., 1997; Golding and Spruston, 1998; Vetter et al., 2001).

#### Consequence 2: Synaptic background activity enhances the propagation of fast dendritic spikes

Interestingly, in this biophysical model, background activity tended to facilitate forward-propagating spikes. The probability of dendritic spike initiation was significantly affected by background activity (Fig. 6B, left, in-vivo–like), and a large fraction (see below) of dendritic spikes could forward-propagate over large distances and reach the soma (Fig. 6C, in-vivo–like), a situation which did not occur in quiescent states. To explain this effect of background activity on dendritic spikes, we compared different background activities with equivalent conductance but different amplitudes of voltage fluctuations. Fig. 6B (right panel) shows that the probability of spike initiation, for fixed stimulation amplitude at 100 μm from soma, was zero in the absence of fluctuations, but steadily raised for increasing fluctuation amplitudes, showing that subthreshold stimuli are occasionally boosted by depolarizing fluctuations. Propagating spikes can also benefit from this boosting to help their propagation all the way up to the soma. The same picture was observed for different morphologies, passive properties and for various densities and kinetics of voltage-dependent currents (Rudolph and Destexhe, 2003): in-vivo–like activity minimally affected backpropagating spikes but always boosted forward-propagating spikes. Thus, under in-vivo–like conditions, even small EPSPs have a chance to initiate a fast dendritic spike, which itself has a chance to propagate and reach the soma. These probabilities can be small, but are never zero.

#### Consequence 3: Synaptic background activity diminish the location-dependence of synaptic efficacy

The computational model was next used to quantitatively evaluate the consequences of forward dendritic spike facilitation in terms of the impact of individual EPSPs at the soma. In quiescent conditions, the model was adjusted to the passive parameters estimated from whole-cell recordings in vitro (Stuart and Spruston, 1998), yielding a significant passive attenuation of synaptic inputs (Fig. 7A). Under in-vivo– like conditions, the model showed a marked increase in voltage attenuation (Fig. 7A, in-vivo–like), which was due to the tonically activated conductance of synaptic background activity. Next, the impact of EPSPs was assessed with active dendrites, by computing the post-stimulus time histogram (PSTH) over long periods of time with repeated stimulation of single, or groups of colocalized excitatory synapses. The PSTHs obtained for stimuli occurring at different distances from the soma (Fig. 7B) show that, in this case, the “efficacy” of these synapses is roughly location independent, as calculated from either the peak (Fig. 7C) or the integral of the PSTH (Fig. 7D). The latter can be interpreted as the probability that a somatic spike is specifically evoked by a synaptic stimulus. Thus, this biophysical model suggests that, under in-vivo–like conditions, there is a severe passive attenuation, but the impact of individual synapses on the soma is nearly independent on their dendritic location, a situation which is often called “synaptic democracy”.

**Figure 7:**
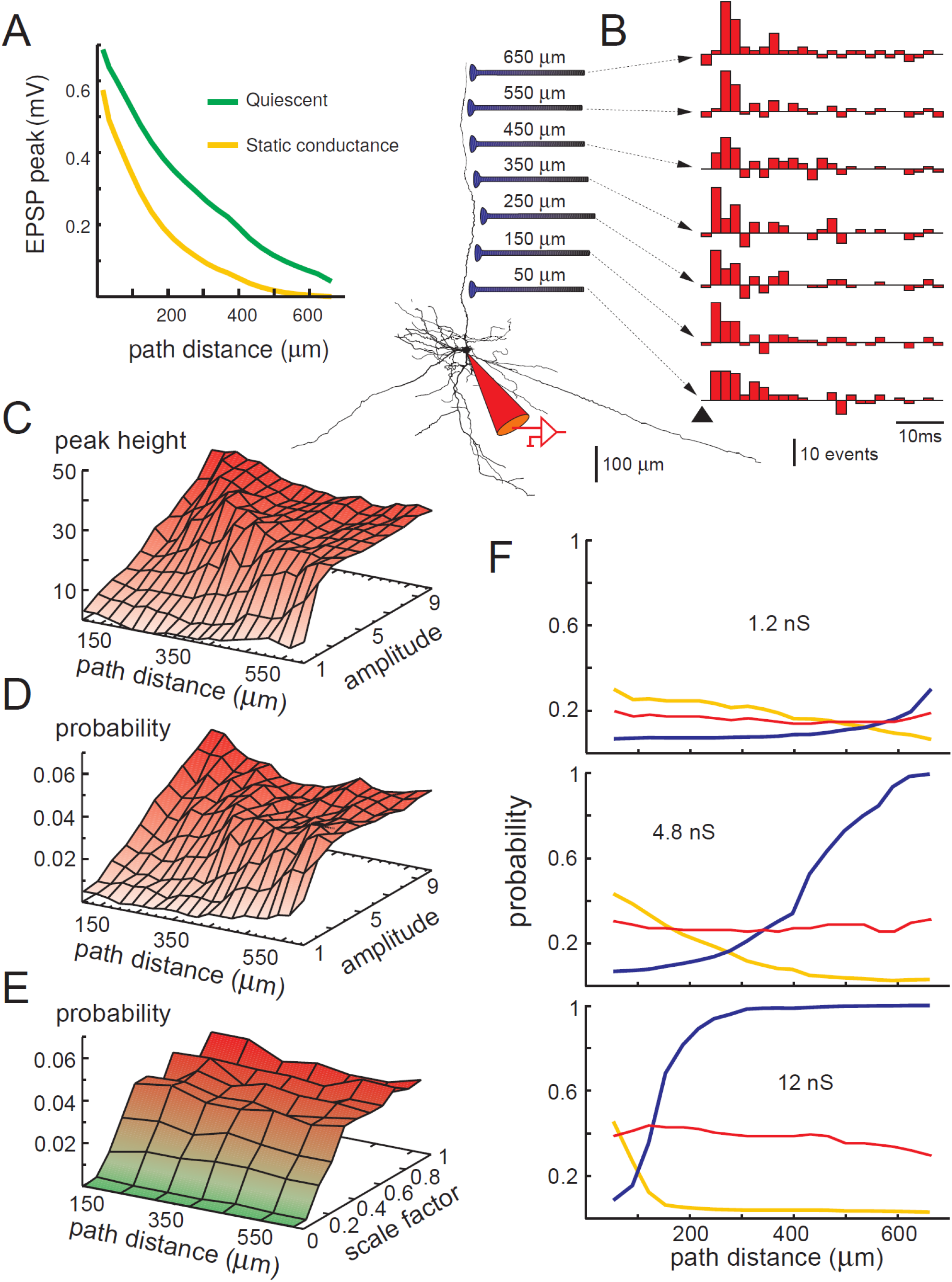
Reduced dependence of the somatic response to the location of synaptic stimulation under in-vivo–like conditions. **A**. Impact of background activity on the attenuation of EPSPs in a passive cortical pyramidal neuron. The neuron simulated in indicated on the right (reconstructed Layer VI pyramidal cell from cat), as well as the location of the synapses tested. The peak of the EPSP is represented as a function of distance to soma. Two sets of passive properties were used, one corresponding to a quiescent neuron (green), and another one with augmented leak conductance to match the conductance due to in-vivo-like activity (yellow). **B**. Post-stimulus time histograms (PSTHs) of responses to identical AMPA-mediated synaptic stimuli (12 nS) at different dendritic locations (cumulated over 1200 trials after subtraction of spikes due to background activity). **C**. Peak of the PSTH as a function of stimulus amplitude (from 1 to 10 co-activated AMPA synapses; conductance range: 1.2 to 12 nS) and distance to soma. **D**. Integrated PSTH (probability that a somatic spike was specifically evoked by the stimulus) as a function of stimulus amplitude and distance to soma. Both C and D show location independence. **E**. Integrated PSTH computed as in D, for various intensities of background activity obtained by scaling the release frequencies at all terminals (stimulus: 12 nS). The plateau region (red) shows that the global efficiency of synapses, and their location independence, are robust to changes in network activity. **F**. Comparison of the probability of evoking a dendritic spike (blue) and the probability that an evoked spike translated into a somatic/axonal spike (orange). Both were represented as a function of the location of the stimulus, for three AMPA-mediated stimulus amplitudes (1.2, 4.8 and 12 nS, respectively). The red curves represent the multiplication of the two other curves (x10), leading to the probability of somatic spike specifically evoked by the stimulus. This probability was nearly location independent for all stimulus amplitudes. Modified from Rudolph and Destexhe, 2003.

To determine the mechanisms underlying this reduced location dependence, we investigated the impact of individual synapses as a function of the intensity of background activity. To do this, we repeated the same stimulation protocols for various background intensities by scaling the release rates at excitatory and inhibitory terminals, which allows one to continuously change from quiescent to in-vivo–like conditions. Computing the cumulated PSTH for subthreshold stimuli showed a near-zero flat trace in quiescent conditions (Fig. 7E, green region). For increasing background intensities, the cumulated PSTH steadily rose, showing that previously subthreshold stimuli evoke detectable responses in the presence of background activity. Interestingly, the response reached a “plateau” where it was nearly independent of both synapse location and background intensity (Fig. 7E, red region). This region corresponds to estimates of background activity based on intracellular recordings in vivo (Pare et al., 1998). Thus, a wide range of background activities seem to equivalently affect how synaptic stimuli evoke somatic spikes, and set the neuron in a mode where the efficacy of individual inputs is nearly location-independent.

To show that this location-independent mode depends on forward-propagating dendritic spikes, we selected, for a given synaptic location, all trials which evoked a somatic spike. These trials represented a small portion of all trials: from 0.4 to 4.5% depending on the location and the strength of the synaptic stimuli. For these “successful” selected trials, the somatic spike was always preceded by a local dendritic spike evoked by the stimulus. In the remaining “unsuccessful” trials, there was a proportion of stimuli (55–97%) which evoked a dendritic spike but failed to evoke somatic spiking. This picture was the same for different stimulation sites: A fraction of stimuli evokes dendritic spikes, and a small fraction of these dendritic spikes successfully evokes a spike at the soma/axon.

We quantified this further as a function of distance to soma. The chance of evoking a dendritic spike was higher for distal stimuli (Fig. 7F, blue), but the chance that a dendritic spike leads to soma/axon spikes was higher for proximal sites (Fig. 7F, orange). Remarkably, these two effects compensated such that the probability of evoking a soma/axon spike was approximately independent on the distance to soma (Fig. 7F, red). This effect was seen for different stimulation intensities (Fig. 7F), and was also found in different pyramidal cell morphologies (Rudolph and Destexhe, 2003).

These results show that the reduced location-dependent impact of synaptic events (synaptic democracy) under in-vivo–like conditions can be explained here from the probabilities of initiation and forward-propagation of fast dendritic spikes. It seems therefore a general phenomenon due to the effect of background activity on local spike initiation and propagation in dendrites, and is probably valid for all cellular morphologies dendrites capable of generating fast spikes. Indeed, the same phenomenon was observed for different reconstructed pyramidal cell morphologies from Layer II-III, Layer V and Layer VI from cat (Rudolph and Destexhe, 2003), so we expect that it would be valid for very large pyramidal cells, such as in human cortex.

## Discussion

In this paper, we have shown that the dendrites of pyramidal neurons generate far more fast local dendritic spikes than somatic spikes, and delineated several possible computational consequences of such local dendritic spike activity.

The experimental observations of the overabundance of dendritic spikes over somatic spikes support early theories of dendritic processing and plasticity. For example, the dendritic spike mediated learning rule (Figure 1 (Mehta, 2004)) relies on the occurrence of plenty of dendritic spikes and depolarization, which is now observed experimentally. In contrast, the standard Hebbian learning rule requires backpropagation of somatic action potential in dendrites, which is highly limited even at the proximal dendrites due to perisomatic inhibition, and nearly impossible in the distal dendrites and in the thin branches. The DAP mediated learning rule can explain recent observations about silent place cells in hippocampus. Here, neurons not only remain silent throughout a maze, but show no subthreshold somatic bump of activity. Yet, neural depolarization causes spiking activity and the establishment of place fields(Lee *et al.*, 2012). This can’t be easily explained by STDP or somatic spike mediated plasticity. But, this can be explained by the existence of a dendritic place field that failed to cause somatic depolarization without current injection, but caused somatic spiking with cellular depolarization.

The dendritic spiking mediated learning can also explain the paradoxical yet well-established observation that place cells turn on abruptly after a few traversals of an environment, even a familiar environment (Mehta McNaughton, B. L. & Bowers, 1997). While the anticipatory dynamics of established place cells can occur via STDP or dendritic learning rule (Mehta *et al.*, 1997, 2000; Mehta, 2015), the abrupt appearance of place cells in the absence of prior somatic spiking is difficult to explain via STDP, but can be readily explained by DAP mediated plasticity. Further, the dendritic location dependent LTP versus LTD has since been observed in proximal versus distal dendrites. Thus, DAP endow neurons with greater computational power and information capacity (Mel, 1993)(London & Häusser, 2005)(Johnston *et al.*, 1996)(Kumar & Mehta, 2011)(Harnett *et al.*, 2012) by turning dendritic branches into computational subunits with branch-specific plasticity(Golding *et al.*, 2002)(Mehta, 2004, 2015). Additionally, the observation that dendrites generate five to ten times as many spikes as the soma during natural behavior suggests that neuronal networks in vivo operate in a regime where the dendritic and somatic activities are decoupled, perhaps via inhibition, thereby greatly enhancing and altering the nature of network computations. Early evidence supports this hypothesis; experiments in virtual reality reveal that the soma and distal dendrites of hippocampal neurons may show very different rhythms, largely mediated by inhibition (Safaryan & Mehta, 2021).

Biophysical models, which were constrained from intracellular measurements of synaptic background activity in vivo, allowed us to look more in detail into the mechanisms underlying dendritic spike initiation and propagation. The biophysical models show that with in-vivo-like activity, the initiation of local dendritic spikes is more likely than in quiescent conditions. This is supported by the recordings of dendrites showing an intense local spiking activity (Moore et al., 2017). A second consequence is that background activity is not necessarily detrimental to dendritic spike propagation. To the contrary, we observed that in conditions where distal spikes normally do not reach the soma (Fig. 6C, quiescent), the presence of background activity enables a small proportion of these spikes to reach the soma (Fig. 6C, in vivo-like). Thus, the probability that a distal synapse speaks to the soma is small, but non-zero in this case (as it would be in a quiescent neuron). Third, we found a remarkable property that the probability of spike initiation and propagation have opposite distance dependence and therefore tend to compensate and lead to a synaptic efficacy that is much less dependent on location (Fig. 7F).

This notion of synaptic democracy was introduced in a previous study (Magee and Cook, 2000), where it was shown that in hippocampal pyramidal cells, there is an increase of the synaptic conductance with distance to soma, which compensates for dendritic attenuation. However, this conductance scaling was not found in pyramidal cells from cerebral cortex (Williams and Stuart, 2002), which suggests that synaptic democracy, if present, should use different mechanisms in cerebral cortex. It was found, similar to a previous proposal (Shepherd et al., 1985), that coincident synaptic releases can trigger local dendritic spikes that can propagate to soma (Williams and Stuart, 2002). We overviewed here a third possible mechanism, which differs from synaptic scaling or coincidence detection, and relies on the presence of synaptic background activity (Rudolph and Destexhe, 2003). According to this mechanism, the efficacy of each synapse is probabilistic and roughly independent on its position with respect to soma. The mechanism works for different dendritic morphologies and does not require that synaptic inputs are co-localized to trigger dendritic spikes, so could in principle be applied to nearly-random connectivity in cerebral cortex.

Interestingly, the fact that this “stochastic” mode of integration applies to different dendritic morphologies (Rudolph and Destexhe, 2003) suggests that this mechanism could be a way to enable the building of very complex networks with several thousand synapses per cell. Such networks would necessarily require neurons to grow extended dendritic morphologies, and yet be able to integrate the information from very distal synapses. It is to be noted that the biophysical models overviewed here only investigated the efficacy of single excitatory inputs. Further models should investigate how combined synaptic inputs (possibly excitatory and inhibitory) are integrated in such stochastic conditions, and how such probabilistic properties impact on more global computations at the network level.

Taken together, experiments and biophysical models suggest that local dendritic spikes implement an “ongoing” spike traffic in the dendrites, which is clearly visible in local dendritic recordings in awake animals. Within this ongoing activity, the efficacy of additional synaptic inputs (as quantified by the probability of evoking a somatic spike) is roughly similar for all synapses. This property is only possible if the excitable dendrites are subject to ongoing nearly-random synaptic background activity, which seems to correspond to the typical asynchronous and irregular activity in the awake brain (Dehghani et al., 2017). Biophysical models therefore predict that pyramidal neurons in vivo should behave probabilistically, through fast dendritic spikes, and that the communication between distal synapses to soma is globally facilitated by background activity. So, similar to stochastic resonance phenomena (Wiesenfeld and Moss, 1995), noise is not a nuisance but participates to enhance information processing, a general theme which also applies to the network level (Zerlaut and Destexhe, 2017).

## Acknowledgements

AD was supported by CNRS and European Community (H2020-945539). MM was supported by NIH grant #1U01MH115746 and AT&T.

